# Characterising the neural time-courses of food attribute representations

**DOI:** 10.1101/2025.05.13.653572

**Authors:** Violet J. Chae, Tijl Grootswagers, Stefan Bode, Daniel Feuerriegel

## Abstract

Dietary decisions involve the consideration of multiple, often conflicting, food attributes that precede the computation of an overall value for a food. The differences in the speed at which attributes are processed play an important role; however, it is unknown whether different attributes are processed over distinct time windows. We mapped the neural time-courses of 12 choice-relevant food attributes. Participants (*N* = 110) viewed food images while we recorded brain activity using electroencephalography (EEG). A separate group of participants (*N* = 421) rated the same images on nutritive properties (healthiness, calorie content, edibility, and level of transformation), hedonic properties (tastiness, willingness to eat, negative and positive valence, and arousal), and familiarity (previous exposure, recognisability, and typicality). Using representational similarity analysis, we quantified differences in patterns of multivariate EEG signals across foods and assessed whether the structure of these differences was correlated with differences in attribute ratings. We observed similar correlation time-courses for many attributes. There was an early window of correlations (∼200 ms from image onset), followed by sustained windows of correlation from ∼400-650 ms. Using principal components analysis, we identified a set of broader constructs that accounted for variance in ratings across multiple attributes, and were also correlated with the EEG data. Our results indicate that food attributes important for choice are represented rapidly and in parallel, over similar time windows. Furthermore, we reveal that broad dimensions underlying individual attributes are also represented in the neural activity with distinct time-courses, indicating a multilevel structure of food attribute representations.

## 1 Introduction

We make dietary decisions frequently throughout each day. These are complex decisions involving the rapid processing of multiple, often conflicting, food attributes. The processing of healthiness, tastiness, and calorie content of foods have been studied extensively in behavioural and neuroimaging studies (Duraisingam et al., 2021; Harris et al., 2013; Lim et al., 2018; Meule et al., 2013; Sullivan et al., 2015; Toepel et al., 2009). However, it is less known how other key features of foods are represented in the brain. Here, we aimed to shed light on the neural representations of a wide range of food attributes that are important for dietary choice.

A growing body of neuroimaging research suggests that dietary decision making involves two processes: the appraisal of the decision option in relation to multiple relevant attributes (e.g., the healthiness and tastiness of a food) and the integration of these representations into an overall value signal that determines one’s choice (e.g., to eat the food or to not eat the food; Hare et al., 2011; Hutcherson et al., 2012). Previous work has shed light on related aspects of food cognition, such as food-related inhibitory control (measured using food go/no-go tasks; e.g., Allen et al., 2022; Carbine et al., 2017), attention (assessed using food oddball tasks; e.g., Babiloni et al., 2009), and associative learning (measured using conditioning paradigms; e.g., Franken et al., 2011). For a methodological review of food-related EEG studies, see Zsoldos et al. (2022). Here, we focus on the appraisal of food options on relevant attributes that occurs prior to the integration of choice-relevant value signals. There are numerous decision-relevant food attributes (Chen & Antonelli, 2020; Schulte-Mecklenbeck et al., 2013), and it is unknown whether these attributes are processed over similar time windows relative to the presentation of a food item. Behavioural evidence indicates that food attributes can differ in how quickly they are integrated into one’s value signals, leading to the over-weighting of attributes thought to be processed more quickly (e.g., tastiness) and the under-weighting of attributes thought to be processed more slowly (e.g., healthiness) in food choice (Lim et al., 2018; Sullivan et al., 2015). To explain these findings, it has been proposed that when presented with a food choice, basic attributes relevant in the short-term (e.g., tastiness) are represented more easily and quickly than abstract attributes relevant in the long-term (e.g., healthiness; Liberman & Trope, 2008). However, the existing neuroimaging evidence is mixed, with some studies reporting neural correlates of healthiness information emerging earlier than those of tastiness (Harris et al., 2018; Schubert, Rosenblatt, et al., 2021).

Food attributes, along with features of other highly biologically relevant stimuli such as faces, animals, and tools, are likely represented in neural signals within several hundred milliseconds after image onset (Carlson et al., 2013; Collins et al., 2018; Grootswagers, Robinson, & Carlson, 2019). An ideal method for examining such rapid neural processes is electroencephalography (EEG), a non-invasive method used to record electrical activity in the brain using electrodes placed on the scalp. EEG has a high temporal resolution that allows one to disambiguate events that occur over short time windows of tens or hundreds of milliseconds. Event-related potentials (ERPs; averaged EEG responses time-locked to an event) can be used to assess the time-courses of neural activity following food image or food cue presentation. Commonly, ERPs evoked by two classes of stimuli (e.g., healthy versus unhealthy foods) are compared. Differences in ERP amplitudes over a given time window have been interpreted as the neural activity in that time window reflecting information about the food attribute (in this case, healthiness of the foods). Previous studies have reported that key food attributes are reflected in both early ERP components (∼100-200 ms after food image onset), thought to be relevant for attentional modulations during early sensory processing, as well as later, more sustained ERP components (∼300-700 ms after food image onset) implicated in higher-level evaluative processes. Multiple studies have found this pattern of results for healthiness (Harris et al., 2013; Rosenblatt et al., 2018), tastiness (Harris et al., 2013; Schubert, Smith, et al., 2021; Schwab et al., 2017), and calorie content (Duraisingam et al., 2021; Meule et al., 2013; Toepel et al., 2009). However, ERP analyses typically focus on signals at a single electrode (or an average of nearby electrodes). Consequently, signals that are distributed across other channels may be missed. Multivariate methods, which assess distributed patterns of neural activity across multiple channels, can address these limitations. Multivariate methods are more sensitive to subtle, distributed differences in neural activity (Hebart & Baker, 2018) and therefore are better suited for examining the fine-grained time-courses of food attribute representations. Using multivariate pattern analysis, one study found that healthiness, tastiness, and willingness to eat information were decodable from patterns of EEG activity from around 500-800 ms after food image onset (Schubert, Rosenblatt, et al., 2021).

As outlined above, most existing work on food choice have focused on healthiness, tastiness, and calorie content as the main attributes that drive choice (e.g., Harris et al., 2013; Schubert, Rosenblatt, et al., 2021; Schubert, Smith, et al., 2021; Sullivan et al., 2015; Toepel et al., 2009). However, there are many other features of foods that may be relevant to dietary decisions. Emerging evidence suggests that the level of transformation (i.e., how much work was required to prepare the food) and edibility are also encoded in the neural activity during food viewing (Coricelli et al., 2019; Moerel et al., 2024). Coricelli et al. (2019) presented participants with images of processed and unprocessed foods matched for calorie content. ERP amplitudes differed as early as 130 ms after food image onset between the two classes of foods, suggesting that similarly to calorie content, level of processing of foods is represented in early neural signals. Familiarity is an important influence on food choice (Blechert et al., 2014; Foroni et al., 2013; Steptoe et al., 1995), but few studies have examined the neural processing of food familiarity in healthy individuals. One study did not observe differences in ERPs evoked by familiar and unfamiliar foods, however they only examined later time windows (500-1000 ms after food image onset; Stuldreher et al., 2023). Whether food familiarity is reflected in the neural signals in earlier time windows (e.g., reflecting early attentional capture for either novel or familiar foods) is yet to be investigated. It is also important to consider different aspects of food familiarity that may differentially influence dietary decisions: frequency of exposure in daily life, recognisability, and how typical the food is of a broader category. Emotion-related attributes such as valence and arousal also differ among foods and likely influence choice (Blechert et al., 2014; Foroni et al., 2013; Schacht et al., 2016). Consistent with findings for other key food attributes, valence is thought to be reflected in early (∼100 ms) and later, more sustained (∼300-700 ms) ERP components (Schacht et al., 2016), but these findings are limited to foods that are positive in valence. Taken together, this evidence suggests that the neural representations of numerous food attributes are important for dietary choice. However, these neural representations are yet to be comprehensively investigated.

One major issue in delineating the neural time-courses of different food attributes is that ratings of many food attributes are strongly correlated. For example, foods perceived to be lower in calories tend to be viewed as healthier, while foods that are thought to be tastier tend to also be judged as more positive in valence. This makes it difficult to determine whether the neural activity is reflecting information about a single food attribute independently of other correlated attributes. Previous work has attempted to decorrelate these food attributes by careful balancing of stimuli (e.g., Schubert, Rosenblatt, et al., 2021) and statistically controlling for effects of correlated attributes (Moerel et al., 2024), however this is difficult when evaluating a broad range of attributes. Another issue is the limited variance in some food attributes within a stimulus set. People tend to perceive foods to be tasty, positive in valence, and desirable for the most part. To increase the variance in these dimensions, some studies have included rotten or mouldy foods, half-eaten foods, or foods that have snails or ants on them (e.g., Piqueras-Fiszman et al., 2014). However, it is possible that these stimuli may not be perceived as foods at all, and instead perceived as inedible (Becker et al., 2016). Few studies have examined foods that are edible but unappetising or unfamiliar. Increasing the variance of these attributes in a stimulus set may improve our ability to map the neural time-courses of these attributes. Lastly, most studies did not control for low- or mid-level visual features (e.g., colour, silhouette) that may correlate with the food attributes of interest and evoke reliable differences in the neural activity. Patterns of EEG responses that correspond to such visual features can often be decoded using multivariate classifiers (Grootswagers et al., 2024; Grootswagers, Robinson, Shatek, et al., 2019). For instance, more green foods may tend to be healthier, and colour can be decoded from EEG responses across a range of latencies from stimulus onset (Grootswagers et al., 2024). Consequently, the effects of colour may produce differences in ERPs or multivariate classification performance between healthy and unhealthy foods.

Recently, Moerel et al. (2024) demonstrated a novel approach for uncovering the time-courses of multiple food attributes using representational similarity analysis (RSA). RSA is a multivariate method that can be used to look for encoded information in neural activity by comparing between neural and behavioural data based on shared structure in their representational dissimilarity matrices (RDMs). Participants viewed food images at rapid presentation rates (∼100 ms per image) while EEG activity was recorded. Food attribute RDMs were constructed using normative ratings from a separate sample of participants (Foroni et al., 2013). Importantly, Moerel et al. (2024) controlled for the effects of low- and mid-level visual features by constructing visual feature RDMs to include in the partial correlation analyses between the neural RDMs and the food attribute RDMs. Level of transformation, immediate edibility, and calorie content of foods were found to be encoded in the patterns of EEG activity within the first second. On the other hand, there was no strong evidence that valence and arousal were encoded in the neural activity.

Here, we replicate and extend on the work by Moerel et al. (2024) using RSA. We examined the neural time-courses of attribute representations when foods are presented for a longer, more naturalistic viewing period (2 s) using a stimulus set that included edible but unappetising foods to increase variance in key food attributes. Furthermore, we assessed novel food attributes that have not been investigated in this way previously, such as familiarity. We recruited a large sample (*N* = 110) to investigate the neural representations of 12 food attributes that broadly span nutritive properties (healthiness, calorie content, edibility, and level of transformation), hedonic properties (tastiness, willingness to eat, negative and positive valence, and arousal) and familiarity (previous exposure, recognisability, and typicality).

We recorded EEG while participants completed a food categorisation task. We carefully designed this task to separate the attribute representation and the integration processes of decisions. Ratings on 12 food attributes were collected from a separate sample of participants (*N* = 421). Using RSA, similarity structures between the EEG activity and the attribute ratings were compared to examine whether, and importantly *when*, food attributes were represented in the neural activity, while controlling for the effects of low- and mid-level visual features. Furthermore, we aimed to disentangle highly correlated food attributes through two complementary approaches. Firstly, we applied principal components analysis (PCA) to identify shared variance dimensions and examined whether these dimensions were represented in the neural activity. Secondly, we assessed whether there were any unique contributions of each food attribute in the EEG data through partial correlation analyses controlling for the effects of all other food attributes.

## 2 Method

Our study involved three parts. In the first session of the main experiment, participants completed a food categorisation task while EEG was recorded and subsequently provided food attribute ratings for each food image. In the second session of the main experiment, participants completed two behavioural food choice tasks and completed questionnaires probing their dietary style and food motivations. Data collected from the second session were not analysed here, and the descriptions of behavioural tasks and the questionnaires are not included. We also recruited a separate sample of participants who rated each food image with respect to the 12 food attributes. This was carried out to create models of the similarity structure between foods for each attribute. We compared EEG activity time-locked to food image presentations with the 12 food attribute models to examine whether and when neural activity covaried with each type of food attribute rating. This study was approved by the Human Research Ethics Committee of the University of Melbourne (HEIC ID 24850). Data processing and analysis code will be available via the Open Science Framework (https://osf.io/w8c9d/) at the time of publication.

### 2.1 Participants

Participants (*N* = 117) were recruited through the University of Melbourne student portal and advertising via posters around the University of Melbourne. Participants indicated that they were at least 18 years old, were fluent in written and spoken English, did not have a history of any eating disorders, were not on a calorie restriction diet, and did not have any dietary restrictions (e.g., for health or religious reasons). Participants were asked to fast for three hours prior to the start of the first and second sessions. At the end of the second session, participants were reimbursed AUD $50. Five participants were excluded as they did not meet the eligibility requirements or did not complete the study. Two additional participants were excluded due to the high incongruence in their continuous healthiness, tastiness, and willingness to eat ratings and their responses in the food categorisation task. The final sample of 110 participants had a mean age of 26.5 years (SD = 8.23, range 18-57 years). 81 participants identified as female, 28 as male, and one as other.

A separate sample of online participants (*N* = 421) were recruited through the University of Melbourne Research Experience Program. Participants indicated that they were at least 18 years old, were fluent in written and spoken English, did not have a history of any eating disorders, were not on a calorie restriction diet, did not have any dietary restrictions, and had not previously taken part in the main experiment. The online sample had a mean age of 19.75 years (SD = 3.08, range 18-53 years). 326 participants identified as female, 92 as male, and three as other. Participants were awarded course credit at the conclusion of the experiment.

### 2.2 Stimuli

The stimuli included 120 colour images of food items presented against a white background (Figure 1A). 91 images were drawn from the Food-pics database (Blechert et al., 2014) selected to represent a wide range of food categories (e.g., vegetables, meats, fruits, snack foods). 29 images were taken from online image hosting websites (e.g., Flickr, Adobe Stock). These were selected to be generally less appetising and less familiar foods to increase the variance in food attribute ratings.

**Figure 1.**
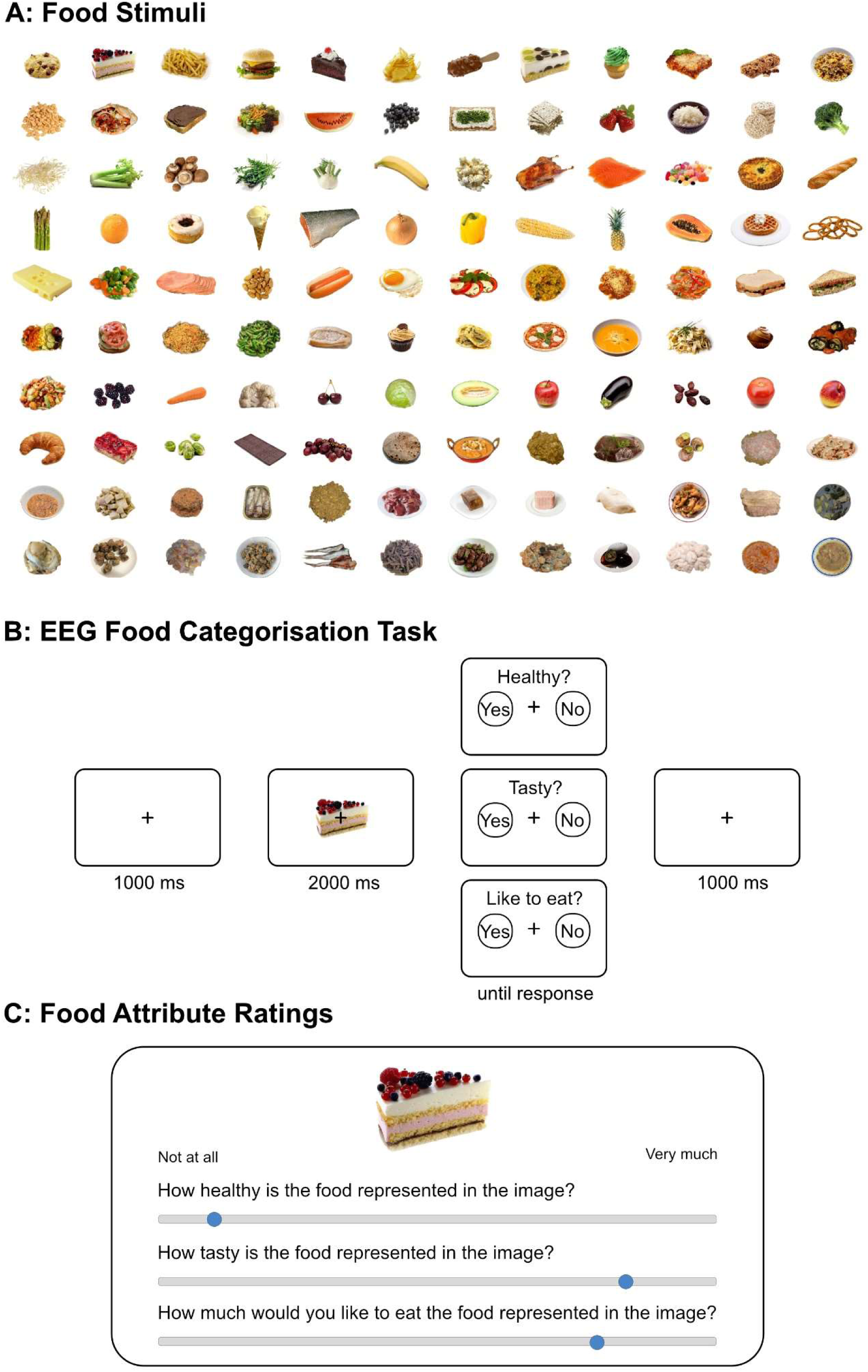
Stimuli and trial diagrams for the EEG and food attribute rating tasks. A) Food stimuli. 120 food images that span a wide range of food categories on a white background. B) EEG Food categorisation task. In each trial, participants were shown a food image for 2 s, followed by a response screen with one of three prompts (‘Healthy?’, ‘Tasty?’, ‘Like to eat?’) and one of two response mappings indicating the corresponding keyboard press for ‘Yes’ and ‘No’ responses (‘z’ key for ‘Yes’ and ‘m’ key for ‘No’, or the reverse). Participants responded as quickly as possible. A fixation cross was displayed between trials for 1 s. C) Food attribute ratings. Participants were shown food images with three food attribute rating prompts at a time, with a sliding scale to indicate their response to each prompt. Ratings on 12 food attributes (healthiness, calorie content, edibility, level of transformation, tastiness, willingness to eat, negative and positive valence, arousal, exposure, recognisability, and typicality) were collected for each food image.

### 2.3 Procedure

During the first session, participants completed a food categorisation task presented using Psychtoolbox (Kleiner et al., 2007) interfacing Matlab R2022a. Afterwards, participants rated each food image on healthiness, tastiness and how much they would like to eat the food (willingness to eat) using Qualtrics.

In the food categorisation task, participants viewed each food item three times: once each across the healthiness, tastiness, and willingness to eat trials while EEG was recorded (Figure lB). Food images were presented centrally, subtending 6.6 × 3.5 degrees of visual angle on a 24.5-inch ASUS ROG Swift PG258Q monitor (1920 × 1080 pixels, 60 Hz refresh rate). Participants’ heads were stabilized using a chin and forehead rest approximately 50 cm from the monitor. Each trial began with a fixation cross presented in isolation for 1 s, followed by the food image with a fixation cross for 2 s. Next, the participants were prompted with “Healthy?” in a healthiness trial, “Tasty?” in a tastiness trial, and “Like to eat?” in a willingness to eat trial, along with a response mapping that indicated which of the left (‘z’) and the right (‘m’) keyboard keys corresponded to ‘Yes’ and ‘No’ responses. There was an equal chance of the left keyboard press indicating a ‘Yes’ response (and the right keyboard press indicating a ‘No’ response) and the reverse for each trial. For each participant, the order of healthiness, tastiness, and willingness to eat trials were randomised. The randomisation of the trial type order and response mapping ensured that participants could not pre-emptively prepare a keypress motor action during the food image presentation period. Participants were asked to respond as quickly and accurately as possible and to pay attention to the switching of the response mapping across trials. Participants completed five practice trials and indicated that they had understood the instructions before starting the main experiment. To reduce eye movement artifacts the participants were instructed to keep their eyes fixated on the fixation cross, which remained on screen throughout the duration of the experimental blocks. Response time was recorded as the time taken from the presentation of the prompt to the participant’s response via keyboard press.

After the food categorisation task, participants rated each food image on healthiness (’How healthy is this food?), tastiness (’How tasty is this food?’) and how willing they were to eat it (’How much would you like to eat this food?’). Participants were asked to use a computer mouse to indicate their response on a sliding scale from ‘Not healthy at all’ to ‘Very healthy’ for the healthiness rating, ‘Not tasty at all’ to ‘Very tasty’ for the tastiness rating, and ‘Not at all’ to ‘Very much’ for the willingness to eat rating. We randomised image presentation order for each participant (while ensuring that the same food image was not presented in two consecutive trials).

In the online rating task, a separate sample of participants indicated their current hunger level (“How hungry are you currently?”) and satiety level (“How full are you currently?”) on a sliding scale from ‘Not at all’ to ‘Very much’. They selected the time elapsed since their last food consumption (“How long ago did you last eat food?”) from one of the following options: ‘Less than an hour ago’, ‘l-3 hours ago’, ‘3+ hours ago’, and ‘I do not know’. Each participant rated a subset of the full stimulus set (40 out of the 120 images) on 12 food attributes, including nutritive properties (healthiness, calorie content, edibility, and level of transformation), hedonic properties (tastiness, willingness to eat, negative and positive valence, and arousal), and familiarity (previous exposure, recognisability, and typicality). It is worth noting that edibility and level of transformation are dissociable: edibility refers to the amount of work *required in the future* to make the food ready to eat from its current state, while level of processing refers to the amount of work *already invested* in the food in its current state. It is possible for a food to be high in edibility but low in level of transformation (e.g., fruits) or conversely, low in edibility and high in level of transformation (e.g., cocoa powder for baking). See Table 1 for the list of food attributes and the corresponding prompts and additional information presented to the participants. We randomised the image presentation order for each participant as well as the order of the food attributes for each image. Participants were asked to indicate their response to each prompt on a sliding scale (0-100) using a computer mouse.

**Table 1.** Food attributes spanning nutritive properties, hedonic properties, and familiarity. Ratings on 12 food attributes were collected using the corresponding prompt for each food image. For edibility, level of transformation, and arousal, additional information was displayed alongside the prompt (in italics).

|  | Food attribute | Prompt |
| --- | --- | --- |
| Nutritive properties | Healthiness | How healthy is the food represented in the image? |
|  | Calorie content | How much calorie content are in the food represented in the image? |
|  | Edibility | How much work WILL BE required to bring the food represented in the image ready to eat?<br>*For example, raw chicken will require more work (i.e., more steps, long time, many ingredients) to get it ready to eat compared to an apple (i.e., less steps, short time, few ingredients).* |
|  | Level of transformation | How much work WAS required to prepare the food represented in the image?<br>*For example, a cake required more work (i.e., more steps, long time, many ingredients) to prepare than rice (i.e., less steps, short time, few ingredients).* |
| Hedonic properties | Tastiness | How tasty is the food represented in the image? |
|  | Willingness to eat | How much would you like to eat the food represented in the image? |
|  | Negative valence | How negatively do you view the food represented in the image? |
|  | Positive valence | How positively do you view the food represented in the image? |
|  | Arousal | How emotionally arousing is the food represented in the image?<br>*Less emotionally arousing foods are calming/relaxing, and more emotionally arousing foods are stimulating/energising.* |
| Familiarity | Exposure | How often do you encounter the food represented in the image? |
|  | Recognisability | How easy is it to identify the food represented in the image? |
|  | Typicality | How typical is the food represented in the image? |

We validated whether the online experiment participants responded in a similar way to the participants in the main experiment. As both samples rated each food image on healthiness, tastiness, and willingness to eat, we compared the average healthiness, tastiness, and willingness to eat rating for each food image between the online and main samples using Pearson’s correlations. Average ratings were strongly correlated for healthiness (*r* = .98), tastiness (*r* = .95), and willingness to eat (*r* = .94), indicating that the two samples appraised the foods similarly on the group level. We therefore opted to use the online sample’s food attribute ratings in our analyses of the neural data from the main sample.

### 2.4 EEG Recording & Data Processing

EEG was recorded using a Biosemi Active II system (Biosemi, The Netherlands) with 64 channels at a sampling rate of 512 Hz using common mode sense and driven right leg electrodes (http://www.biosemi.com/faq/cms&drl.htm). We attached 64 electrodes to the cap according to the international 10-20 system. We also attached eight additional electrodes: two electrodes placed 1 cm from the outer canthi of each eye, four electrodes placed above and below the centre of each eye, and two electrodes placed above the left and right mastoids. Electrode offsets were kept within the range of ±20 *µ*V during recording.

EEG data was processed using EEGLab v2022.1 (Delorme & Makeig, 2004) interfacing Matlab (R2022a). Excessively noisy channels were identified through visual inspection (mean number of bad channels = 0.26, range 0–5) and excluded from average reference calculations and independent components analysis (ICA). Excessively noisy sections or those with large amplitude artifacts were identified through visual inspection and removed. The data were referenced to the average of all channels (excluding excessively noisy channels). One channel (AFz) was removed to compensate for the rank deficiency due to average referencing. EEG data were low-pass filtered at 30 Hz (EEGLab Basic Finite Impulse Response Filter New, default settings). A copy of the dataset was created for the purpose of ICA. A 0.1 Hz high-pass filter (EEGLab Basic Finite Impulse Response Filter New, default settings) was applied to the copied dataset to improve stationarity for the ICA. We used RunICA extended algorithm (Jung et al., 2000) to perform the ICA. The resulting independent component information was copied to the original dataset (e.g., as done by den Ouden et al., 2023). Independent components associated with eye blinks and saccades were identified and removed in line with guidelines in (Chaumon et al., 2015). Next, we interpolated the excessively noisy channels and AFz using spherical spline interpolation.

EEG data were segmented from −100 ms to 1000 ms relative to food image onset. Segments were baseline-corrected using the data from the 100 ms time window prior to each food image presentation. Segments from trials where the participant took longer than 5 s to respond or those containing amplitudes exceeding ±150 *µ*V at any channel were excluded from the analyses. We chose ±150 *µ*V as the amplitude threshold, as RSA is unlikely to be biased by large amplitude signals compared to ERP analyses, in which trial-averaged ERP measures are more susceptible to large artifacts. On average, 346 out of 360 trials were retained (range 283 – 360).

### 2.5 Representational Similarity Analysis

We analysed the data using RSA (Kriegeskorte et al., 2008; Kriegeskorte & Kievit, 2013). Compared to traditional ERP analysis, RSA had distinct advantages given that our research question focussed on characterising the neural time-courses of representations for a broad array of food attributes. This type of analysis allowed for a much more fine-grained examination of the time-points at which the food attribute RDMs were correlated with the neural activity. Furthermore, we could control for the influence of low- and mid-level visual features using partial correlation analyses. For each participant, we created time-varying representational dissimilarity matrices (RDMs; Grootswagers et al., 2017) that characterised the dissimilarity in the neural activity recorded while viewing the food images, for all possible pairs of food images. Next, we created models for the food attributes using the ratings from the online sample. For each food attribute, we created RDMs that characterised the dissimilarity in the group average food attribute ratings for all possible pairs of food images. As low- and mid-level visual features of the images, such as colour and shape, may correlate with the food attributes, we sought to control for the effects of such image features. For example, the colour green may be positively correlated with the healthiness of foods and negatively correlated with calorie content. We created RDMs that encoded the similarity structure regarding such low- and mid-level visual features between our food stimuli. Using partial correlation analyses, we accounted for patterns in the EEG activity that were explained by similarity in the visual features across the stimuli before assessing covariation between EEG signals and the food attributes. This allowed us to compare the neural RDMs to each of the food attribute RDMs to examine whether and when the food attribute information was encoded in the EEG activity, while controlling for the effects of the visual features.

#### 2.5.1 Constructing Representational Dissimilarity Matrices

##### 2.5.1.1 Neural RDMs

We created time-varying neural RDMs for individual participants. Each food image was presented three times to the participants during the food categorisation task, yielding three segments of EEG activity during food image presentation. After the EEG data processing steps described above, which involved rejection of trials with excessively noisy data, between zero to three segments remained for each food image. If no segments remained after trial rejection for a food image, the food image was excluded from further analyses for that participant. For each of the remaining food images, we took the average of the channel amplitudes across the retained segments for each of the 64 channels. For all pairwise combinations of food images, we computed the dissimilarity between the averaged channel amplitudes for all 64 channels at each time-point of the segment. Dissimilarity for each food image pair was measured as 1 minus the correlation coefficient, with 0 indicating the lowest dissimilarity (i.e., highest similarity) and 2 indicating the highest dissimilarity (i.e., lowest similarity). We repeated this process for data at each time-point relative to food image onset to create time-varying RDMs for each participant.

##### 2.5.1.2 Food attribute RDMs

To create the food attribute RDMs, we firstly calculated the group-average food attribute rating for each food image using the data collected during the online rating task. Group-average ratings were rank transformed for each food attribute across the 120 images. We calculated the dissimilarities as the Euclidean distance between the rank transformed group-average ratings for all pairwise combinations of food images to create a rating-based RDM for each food attribute: healthiness, calorie content, edibility, level of transformation, tastiness, willingness to eat, negative and positive valence, arousal, exposure, recognisability, and typicality. A visualisation of this process is displayed in Figure 2C.

**Figure 2.**
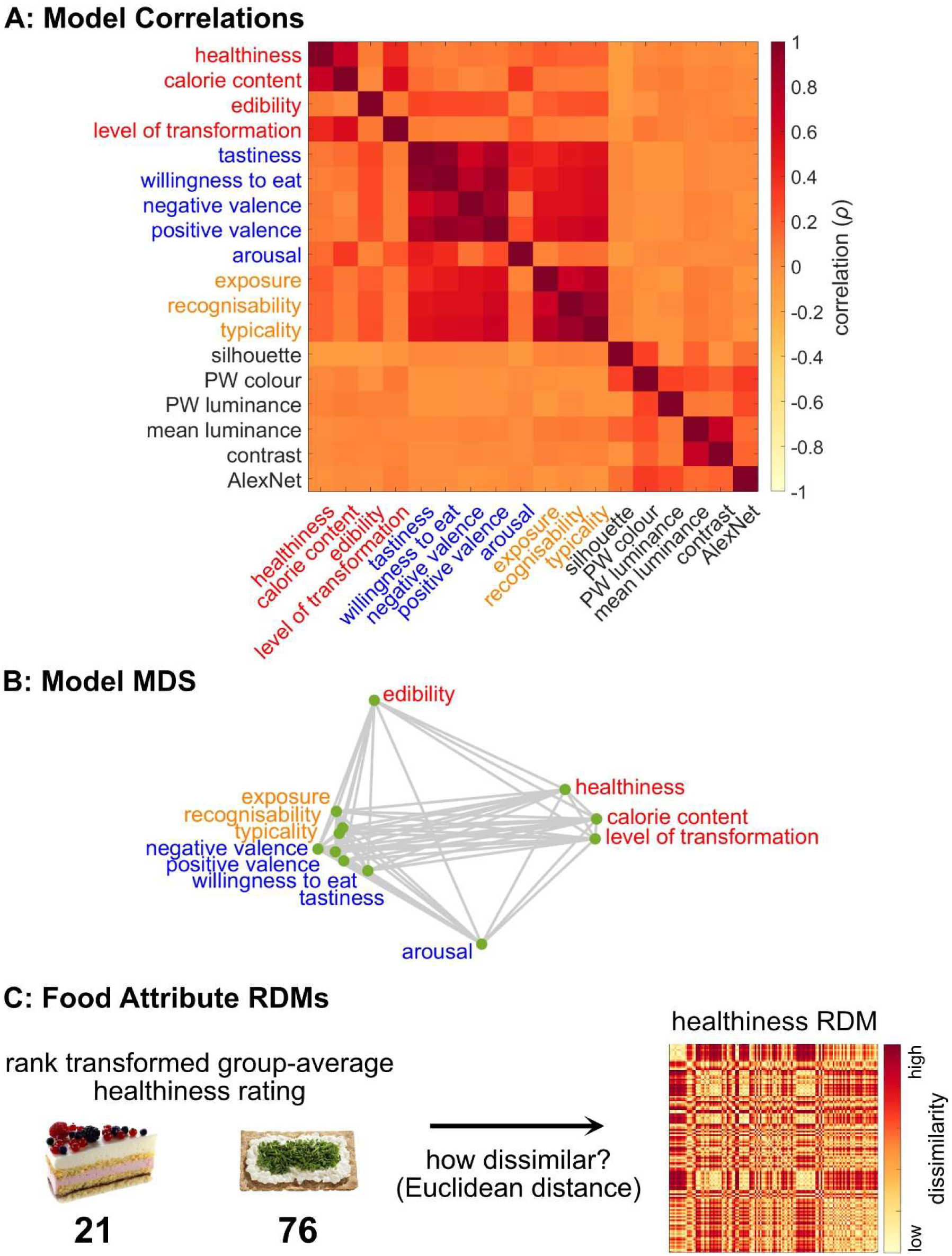
Models used in the representational similarity analysis. A) Model correlations. Correlations between 12 food attribute RDMs and six visual feature RDMs. The food attributes included nutritive properties, hedonic properties, and familiarity. Red indicates positive correlations (i.e., higher similarity) and yellow indicates negative correlations (i.e., lower similarity) between models. B) Model multidimensional scaling. Model correlations between the 12 food attribute RDMs depicted in 2-dimensional space. Smaller distances between the models indicate stronger correlation. C) Food attribute RDMs. Food attribute RDMs were created by calculating the dissimilarity (as measured by Euclidean distance) between the rank transformed group-average attribute rating between all pairwise combinations of food images. For example, we calculated the dissimilarity between the rank transformed group-average healthiness ratings for all possible food pairs (e.g., 21 vs. 76) to construct the healthiness RDM.

##### 2.5.1.3 Visual feature RDMs

Six visual feature RDMs were created to model the low- and mid-level visual features of the food images. The silhouette RDM used vectorized images and measured dissimilarity as the Jaccard distance between binary alpha layers for all possible pairwise image combinations. The pixel-wise colour RDM was based on CIELAB colour values of vectorized images, using correlation as the distance measure. Similarly, the pixel-wise luminance RDM used greyscale values of vectorized images, with correlation as the distance measure. For these conversions, we used the rgb21ab function in Matlab, assuming a D65 white point and an sRGB colour space. The mean luminance RDM calculated dissimilarity across the entire image using the mean greyscale values of the stimuli, with the mean difference as the distance measure. The root mean square contrast RDM was based on the standard deviation of greyscale values across the entire image, also using the mean difference as the distance measure. Lastly, we used a pretrained neural network to model visual similarity (AlexNet; Krizhevsky et al., 2012) in order to control for image features such as shape and complexity. We extracted activations at layer fc8 (the final fully connected layer) and calculated the Euclidean distance for each image pair.

##### 2.5.1.4 Similarities among food attribute and visual feature RDMs

We assessed the similarities among the food attribute and visual feature RDMs by running pairwise correlation analyses for all pairwise combinations of the RDMs (Figure 2A). The similarities between the food attribute RDMs are visualised in the model multidimensional scaling plot (Figure 2B). We included this to examine whether and which food attributes were strongly correlated, as attribute correlation can make it difficult to infer whether the neural activity reflects information about a single food attribute independently of other correlated attributes. We were also interested in examining whether any visual feature RDMs were correlated with the food attribute RDMs, which may indicate a potential confounding effect of image properties that drive visual effects. Tastiness, willingness to eat, negative and positive valence, exposure, recognisability, and typicality RDMs were found to be positively correlated with each other. Healthiness, calorie content, and level of transformation RDMs were found to be positively correlated with each other. From the visual feature RDMs, mean luminance and contrast RDMs were found to be positively correlated with each other.

#### 2.5.2 Comparing Representational Dissimilarity Matrices

We compared similarities over time between neural RDMs and each food attribute RDM using partial correlation analyses, controlling for the effects of visual features. Using correlations to compare RDMs across modalities (e.g., neural, behavioural) is considered standard practice for RSA (Grootswagers et al., 2017; Kriegeskorte et al., 2008; Kriegeskorte & Kievit, 2013) and has been widely used (e.g., Cichy et al., 2014; Grootswagers et al., 2024; Grootswagers, Robinson, Shatek, et al., 2019; Moerel et al., 2024). We first extracted the partial correlation coefficients for each participant between each food attribute RDM and their neural RDMs at each time-point relative to food image onset, controlling for the effects of six visual feature RDMs. Correlations between the visual feature RDMs and the neural RDMs are presented in the Supplementary Figure l.

##### 2.5.2.1 Statistical analyses

To assess whether the neural data and the food attribute RDMs were correlated at a level above chance at the group level, we used a combination of Bayesian and frequentist analyses. Taking the partial correlation coefficients between the food attribute RDMs and each participant’s neural RDMs, we computed Bayes factors using Bayesian one-sample t-tests to determine evidence for above-chance positive partial correlations (i.e., correlation coefficient above 0) at each time-point. For the alternative hypothesis of above-chance positive partial correlations, a half-Cauchy prior was used with a scale factor of .707 (Teichmann et al., 2022). Bayes factors greater than 1 indicate evidence preferentially supporting the alternative hypothesis (positive partial correlation), while Bayes factors less than 1 indicate evidence preferentially supporting the null hypothesis (no positive partial correlation). Specifically, we considered Bayes factors greater than 10 as substantial evidence for the alternative hypothesis, and Bayes factors less than 1/10 as substantial evidence in favour of the null hypothesis (Jeffreys, 1998; Wetzels et al., 2011).

Additionally, we used cluster-based permutation tests (Maris & Oostenveld, 2007; 10,000 permutation samples, cluster forming alpha = .01, family-wise alpha = .05) using functions from the Decision Decoding Toolbox v1.1.5 (Bode et al., 2019) to correct for multiple tests across analysis time windows. For each permutation sample, the sign of the correlation coefficient were flipped for a subset of participants. One-sample t-tests (one-tailed) were then conducted on this permuted data at each time-point. Clusters were formed from sets of adjacent t-values corresponding to *p*-values less than .01. The sum of t-values within each cluster was defined as the cluster’s “mass”. The largest cluster masses from each of the 10,000 permutation samples were used to estimate the null distribution. The cluster masses from the original dataset, where correlation coefficients were not swapped, were then compared to this null distribution. The percentile rank of each cluster mass relative to the null distribution was used to calculate the *p*-value for each cluster. This approach controls for the weak family-wise error rate while retaining sensitivity to detect effects in temporally autocorrelated EEG data (Maris & Oostenveld, 2007).

In the current study, we considered time-points that were statistically significant after application of the cluster-based permutation test and at which Bayes factors exceeded 10 as evidence for a positive partial correlation.

##### 2.5.2.2 Similarity among the time-courses of correlations

We used dynamic time warping to measure the similarity between the time-courses of correlations with the neural RDMs for each of the food attributes. Dynamic time warping is an algorithm that measures the similarity between two temporal sequences, even if they vary in speed or length. This method is shape-based and is therefore suitable for comparing time series data that may not be perfectly aligned temporally (Aghabozorgi et al., 2015). Previous work has used this method to quantify similarity between time-courses of representations derived using RSA (Teichmann et al., 2024). We used this method to measure the distance (Euclidean) between the correlations over time for all pairwise combinations of food attributes. Smaller distance indicates more similar time-courses of correlations between the food attributes and the neural RDMs.

### 2.6 Investigating covariation among food attributes

We observed that many food attributes covaried with each other (Figure 2A, 2B) which likely contributed to the similarity in the time-courses of correlations with the EEG data (Figure 5). To shed light on this, we took two complementary approaches.

#### 2.6.1 Capturing broad underlying dimensions

We performed PCA using the group-average ratings for each of the food images for all 12 food attributes. The rating data was rank transformed for consistency with the food attribute RDMs and then standardised (*M* = 0, SD = l) for each attribute prior to extracting components. Components with eigenvalues greater than 1 were retained (Kaiser’s rule; Kaiser, 1960). The final solution retained two components, which explained 85.5% of the total variance.

We created principal component RDMs for each of the first two components. We used the Euclidean distance between the component loadings for all pairwise combinations of food images as the dissimilarity measure. To assess whether the underlying dimensions captured by the principal components were represented in the EEG signals, each principal component RDM was correlated with each participant’s neural RDMs at each time-point, using partial correlation analyses to control for the six visual feature RDMs. We tested for above-chance positive partial correlations at the group level using Bayesian and frequentist t-tests as described above.

#### 2.6.2 Unique contributions to the correlations between the neural RDMs and the food attribute RDMs

To assess whether any of the food attributes explained any variance in the neural data not accounted for by other food attributes or visual features, we performed partial correlation analyses to measure the correlation between each food attribute RDM and the neural RDMs controlling for all other food attributes and the visual features. For example, neural RDMs were correlated with the healthiness RDM for each time-point and for each participant, using partial correlations to control for the other ll food attribute RDMs and the six visual feature RDMs. We tested for above-chance positive partial correlations at the group level using Bayesian and frequentist t-tests as described above.

## 3 Results

We examined the neural time-courses of food attribute representations during the appraisal of foods. We used representational similarity analysis (RSA; Kriegeskorte et al., 2008; Kriegeskorte & Kievit, 2013) to create models of the EEG activity and food attributes. We used partial correlation analyses to determine the time-points at which the EEG signals and the food attribute ratings were correlated, while controlling for the low- and mid-level visual features. To further our understanding of the covariation between food attributes, we took two different approaches. Firstly, we performed PCA to examine whether the covariation between ratings could be captured in components, and whether these components also covaried with the EEG data. Furthermore, we used partial correlation analyses to determine whether there were time-points at which the food attributes were uniquely correlated with the EEG activity after controlling for the rest of the food attributes and the visual features.

### 3.1 Time-courses of food attribute representations

We examined the correlations between the neural RDMs and the food attribute RDMs at each time-point, using partial correlation analyses to control for the six visual feature RDMs. Positive correlations between the neural RDMs and the food attribute RDMs indicate that the representational geometries, estimated by the dissimilarities between all possible food pairs, were similar on the neural level and on the level of food attribute judgements. This can be interpreted as the neural patterns reflecting food attribute information. The strength of the correlation indicates how similar the neural RDMs and the food attribute were, with peaks in correlations indicating the time at which there was the highest level of similarity.

We found that for many food attributes, there was an early peak in correlations with the neural RDMs around 200 ms from food image onset and a later, more sustained window of correlations starting from around 400 to 650 ms onwards. Taking healthiness as an example (Figure 3A), the top plot shows the correlation between the neural RDMs and the healthiness RDM over time, with 0 ms indicating the time of the food image onset. Positive correlation indicates that the neural RDM is similar to the healthiness RDM, which indicates that healthiness information is present in the neural activity at that time.

**Figure 3.**
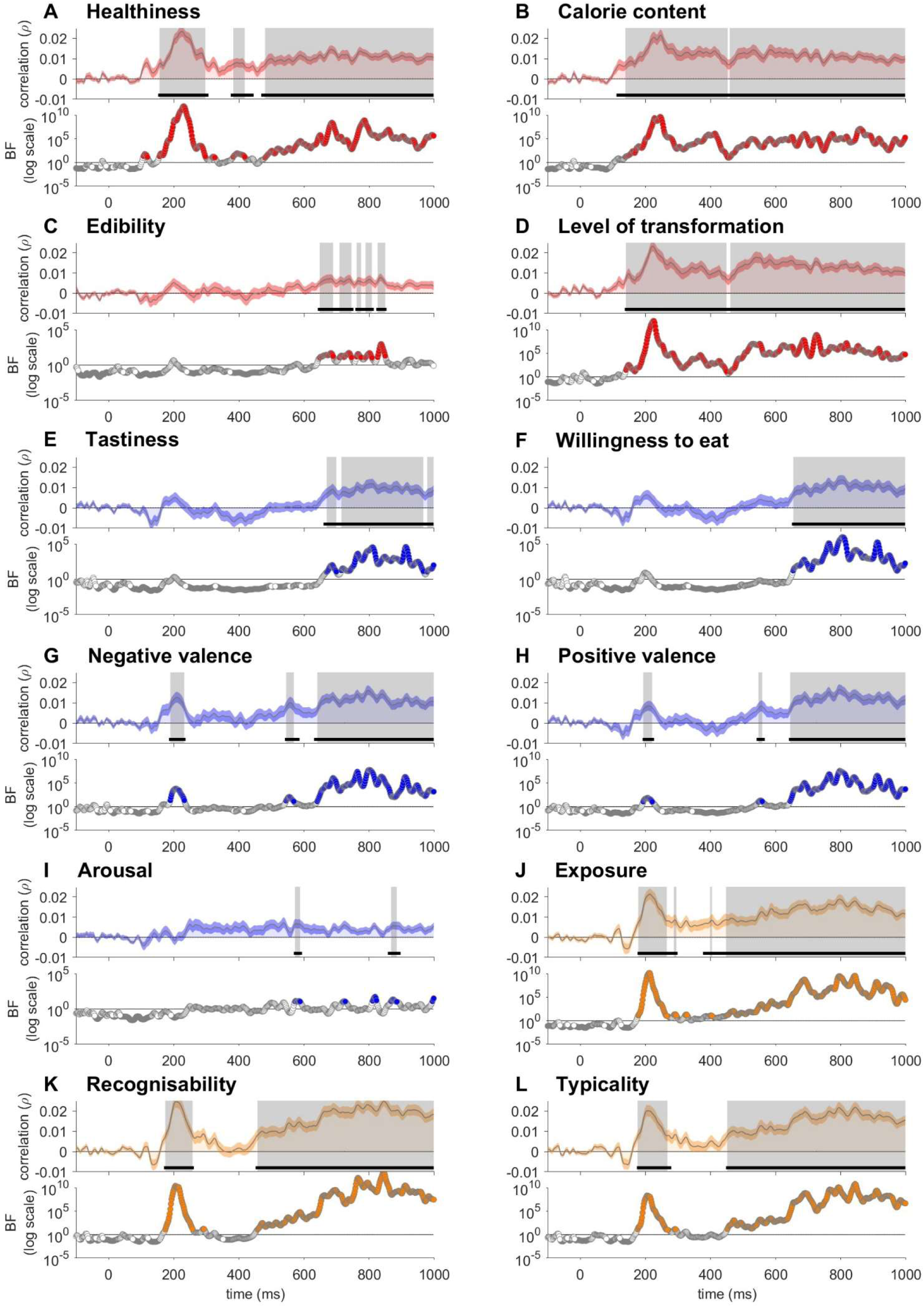
Correlations between the EEG and the food attributes while controlling for visual features. Partial correlations between the 12 food attribute RDMs and the neural RDMs, controlling for the six visual feature RDMs. Nutritive properties (A-D) include healthiness, calorie content, edibility, and level of transformation. Hedonic properties (E-I) include tastiness, willingness to eat, negative valence, positive valence, and arousal. Familiarity attributes (J-L) include exposure, recognisability, and typicality. The shaded area around the plot lines shows the standard error of the mean. The horizontal black lines indicate time windows with statistically significant above-zero correlations after applying multiple comparisons corrections. The coloured circles indicate the time-points at which the BFs were greater than 10, while grey circles indicate the time-points at which the BFs were less than 1/10, and the white circles indicate the time-points at which the BFs were between 1/10 and 10. The grey shaded areas indicate the time windows at which the BFs were greater than 10 and there were statistically significant above-zero correlations after applying multiple comparisons corrections.

#### 3.1.1 Nutritive properties

The correlations for healthiness, calorie content, and level of transformation displayed an early peak (at 227 ms, 248 ms, and 223 ms respectively; Figure 3A, 3B, 3D). For calorie content and level of transformation, there was one sustained window of correlations (starting from 140 ms and 137 ms respectively) that lasted until the end of the analysed time range. Healthiness was found to be correlated from around 152 ms and in later, sustained windows starting from 375 ms and lasting until the end of the analysed time range. Edibility was found to be correlated later than the other nutritive properties, from 648 ms (Figure 3C). Nutritive properties, with the exception of edibility, appeared to be correlated with the early EEG signals (from 100-150 ms) in a sustained manner throughout the analysed time range, with a peak (indicating the time at which there was the highest similarity) around 200-250 ms.

#### 3.1.2 Hedonic properties

The valence attributes were found to be correlated with the neural data from 189 ms for negative valence and 193 ms for positive valence, with an early peak at 209 ms for both attributes (Figure 3G, 3H). Additionally, the negative and positive valence attributes were correlated in later, more sustained windows from 545 ms and 640 ms respectively and lasting until the end of the analysed time range. Tastiness and willingness to eat were correlated later, from 670 ms and 654 ms respectively and sustained until the end of the analysed time range (Figure 3D, 3E). For arousal, there were some time-points at which the BFs exceeded 10 and the cluster-based permutation test for multiple comparisons returned significant results; however, these windows were very small (Figure 3I). Hedonic properties, with the exception of arousal, appeared to be correlated with the later neural activity (from 600-650 ms) in a sustained manner until the end of the analysed time range. For the valence attributes, there was an additional small window of correlations around 200 ms.

#### 3.1.3 Familiarity

Exposure, recognisability, and typicality displayed an early peak at 213 ms, 207 ms, and 205 ms respectively, and a later, more sustained window of encoding starting from 377 ms, 457 ms, and 451 ms respectively and lasting until the end of the analysed time range (Figure 3J-L). Familiarity attributes displayed the highest similarity with the EEG activity around 200 ms, as indicated by the peaks in correlations, and were also correlated with later EEG activity (from 350-450 ms) in a sustained manner until the end of the analysed time range.

#### 3.1.4 Similarities among time-courses

As many of the food attributes shared similarities in their time-courses of encoding, we used dynamic time warping to measure these similarities. In general, attributes within groups were found to share similar time-courses of encoding, as evidenced by smaller distances between the time-courses of correlations (Figure 5). Visual inspection of the plot revealed distinct clusters of attributes that were more similar in their time-courses of correlations, as evidenced by the clustering of yellow cells. Note that we did not use a specific threshold to distinguish high from lower similarity, as the absolute Euclidean distance values are not individually informative. Instead, we used the colour gradients to describe the similarity structure across all pairwise combinations of the correlation time-courses. Out of the nutritive attributes, healthiness and calorie content, as well as calorie content and level of transformation, were found to be similar in their time-courses of encoding. Encoding of many of the hedonic properties (tastiness, willingness to eat, negative and positive valence, but excluding arousal) were also found to be similar over time. Furthermore, the time-courses of encoding for all three familiarity attributes (exposure, recognisability, and typicality) were found be similar. Interestingly, time-courses of encoding for the majority of the hedonic properties (tastiness, willingness to eat, negative and positive valence) were found to be similar to that for familiarity (recognisability and typicality, and exposure). The model correlation matrix (Figure 2A) and the dynamic time warping matrix (Figure 5) were found to be strongly (negatively) correlated (Spearman’s rho = -.94), indicating that the two matrices are structurally similar.

### 3.2 Identifying broad underlying dimensions

We used PCA to identify broad underlying dimensions that explained substantial amounts of variance in food attribute ratings. To assess whether the information captured in these principal components were present in the neural activity, we examined the correlations between each of the principal component RDMs and the neural RDMs over time, using partial correlation analyses to control for the six visual feature RDMs.

#### 3.2.1 PCA results

Eigenvalues were used to determine the number of components to retain. According to Kaiser’s rule (eigenvalues > l), two components were extracted. The first principal component had an eigenvalue of 7.01, explaining 58.4% of the variance, while the second component had an eigenvalue of 3.26, explaining 27.2% of the variance. Together, these two components accounted for 85.5% of the total variance. Factor loadings are presented in Table 2. The first component had large positive loadings for tastiness, willingness to eat, positive valence, and the familiarity attributes (exposure, recognisability, typicality), and a large negative loading for negative valence. The first component also had a negative loading for edibility (for this attribute, low scores indicate high edibility and high scores indicate low edibility). This component likely reflects how appetising the food is, and also maps onto the largest cluster in the model multidimensional scaling plot (Figure 2B). The second component had large positive loadings for calorie content, level of transformation, and arousal, and a large negative loading for healthiness. This component likely taps into how processed the food is, and maps onto the second largest cluster in the model multidimensional scaling plot (Figure 2B).

**Table 2.** Factor loadings for principal components 1 and 2 based on principal component analysis performed with food attribute ratings. Loadings above .3 or below - .3 are deemed large loadings and are in boldface.

| Food attribute | Factor loadings |  |
| --- | --- | --- |
|  | PC1 | PC2 |
| Tastiness | **0.352** | 0.160 |
| Willingness to eat | **0.360** | 0.109 |
| Negative valence | **-0.361** | 0.036 |
| Positive valence | **0.368** | 0.023 |
| Exposure | **0.340** | -0.114 |
| Recognisability | **0.349** | -0.032 |
| Typicality | **0.358** | -0.032 |
| Edibility | -0.248 | 0.002 |
| Arousal | 0.182 | **0.418** |
| Healthiness | 0.128 | **-0.489** |
| Calorie content | -0.024 | **0.541** |
| Level of transformation | -0.059 | **0.489** |

#### 3.2.2 Broad underlying dimensions

The first and second principal component RDMs were correlated with neural signals and showed distinctive time-courses (Figure 4). Both components displayed an early peak at 215 ms and 248 ms respectively. For the first component, this was followed by a sustained window of correlations starting from 642 ms. For the second principal component, correlations were sustained across a large window from 141 ms until the end of the analysed time range. We examined the similarities in the time-courses of encoding between the principal components and the individual food attributes using distance time warping (Figure 5). The first principal component shared similar time-courses of encoding with the attributes that relate to how appetising the food is (tastiness, willingness to eat, valence, exposure, recognisability, and typicality), while the second principal component shared similar time-courses of encoding with the attributes that relate to how processed the food is (healthiness, calorie content, level of transformation, and arousal).

**Figure 4.**
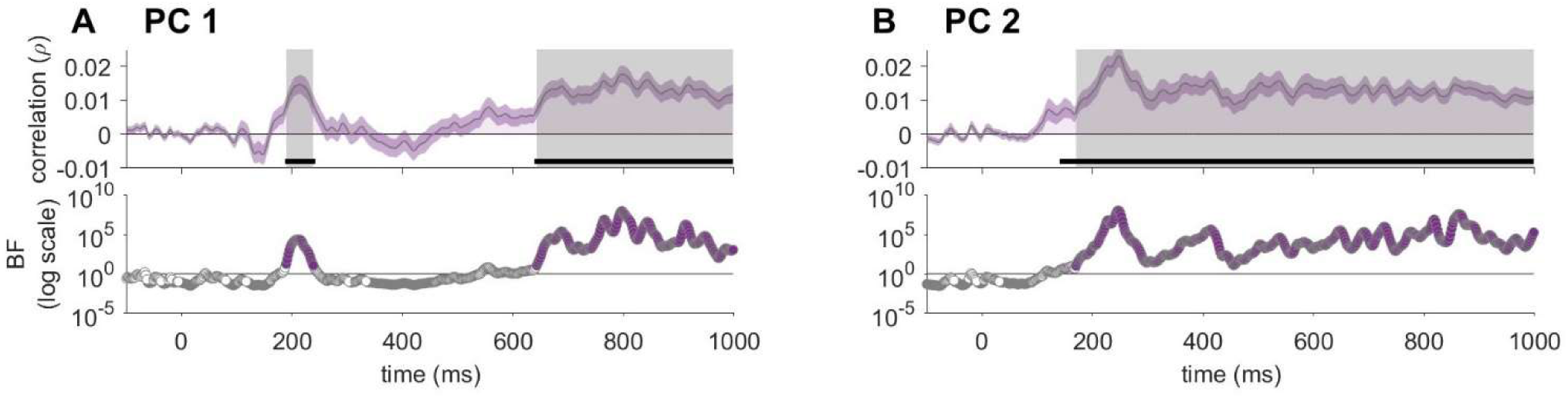
Correlations between the EEG and the principal components while controlling for visual features. Partial correlations between the first (A) and second (B) principal component RDMs and the neural RDMs, controlling for the six visual feature RDMs. The shaded area around the plot lines shows the standard error of the mean. The horizontal black lines indicate time windows with statistically significant above-zero correlations after applying multiple comparisons corrections. The coloured circles indicate the time-points at which the BFs were greater than 10, while grey circles indicate the time-points at which the BFs were less than 1/10, and the white circles indicate the time-points at which the BFs were between 1/10 and 10. The grey shaded areas indicate the time windows at which the BFs were greater than 10 and there were statistically significant above-zero correlations after applying multiple comparisons corrections.

**Figure 5.**
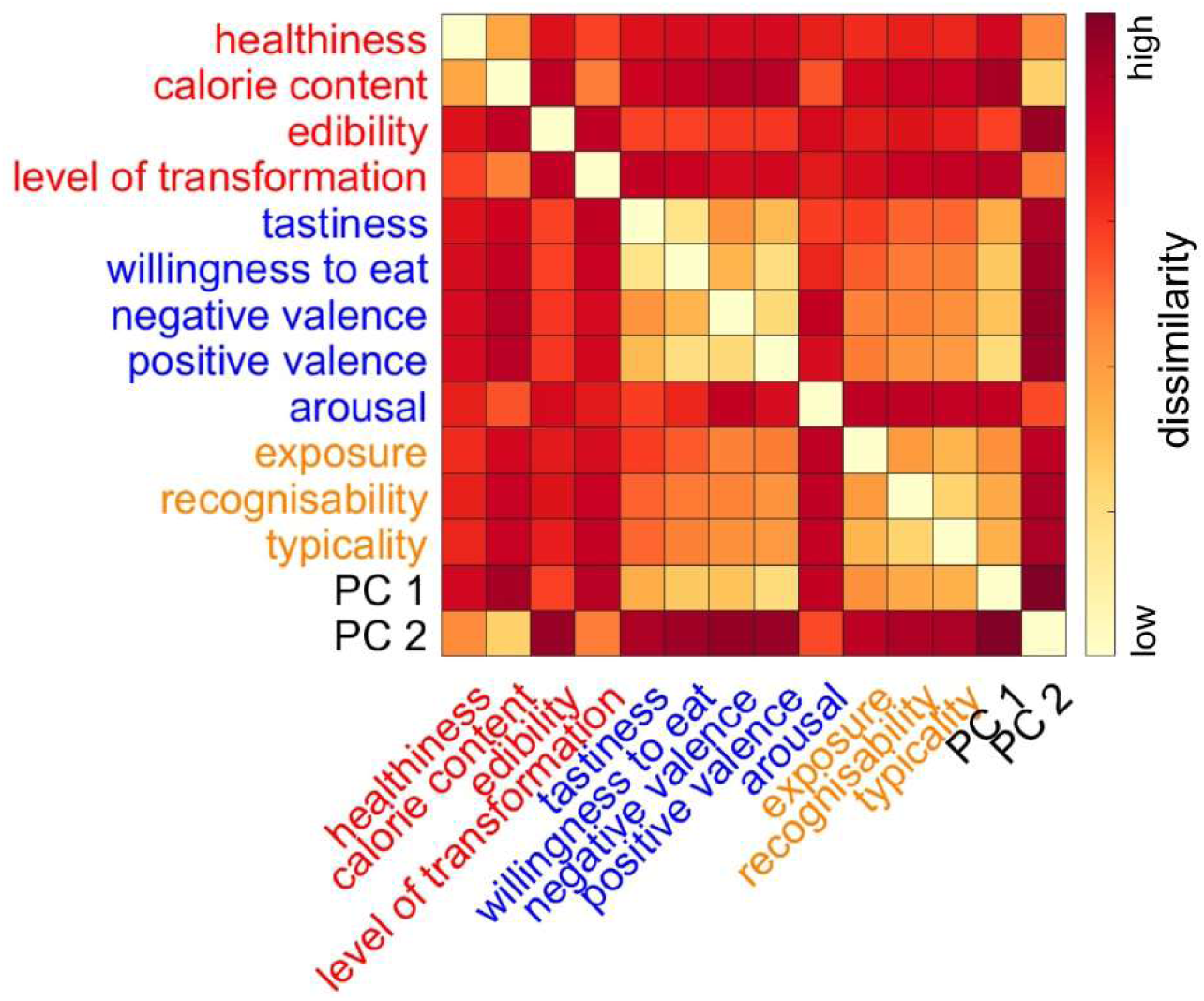
Dynamic time warping to compare similarities between the neural time-courses of food attribute representations. Dissimilarity (measured in Euclidean distance) between the partial correlations over time between the neural RDMs and each of the food attributes, controlling for visual features. Cell colour indicates dissimilarity, with yellow cells indicating lower dissimilarity and red cells indicating higher dissimilarity.

### 3.3 Unique contributions to the variance in neural activity

We next examined whether any food attributes accounted for any variance in the neural data outside of the shared variance explained by other food attributes or visual features. We performed partial correlations with the neural RDMs and each of the food attribute RDMs while controlling for the other ll food attribute RDMs and the visual feature RDMs. This allowed us to evaluate the unique variance in EEG data explained by each food attribute while controlling for the influence of all other food attributes, as well as the visual features. We found that a subset of food attributes across nutritive properties (healthiness, calorie content, and level of transformation), hedonic properties (negative valence and arousal), and familiarity (recognisability) explained unique variance in the neural data.

#### 3.3.1 Nutritive properties

Healthiness, calorie content, and level of transformation explained variance in the neural data after controlling for all other food attributes and visual features (Figure 6A, 6B, 6D). All three attributes were found to be correlated early (from 195 ms for healthiness, 24 ms for calorie content, and 197 ms for level of transformation). Encoding of healthiness and level of transformation displayed an early peak around 203 ms and 221 ms respectively. For level of transformation, there was an additional later window of correlations starting from around 502 ms onwards that was sustained until the end of the analysed time range.

**Figure 6.**
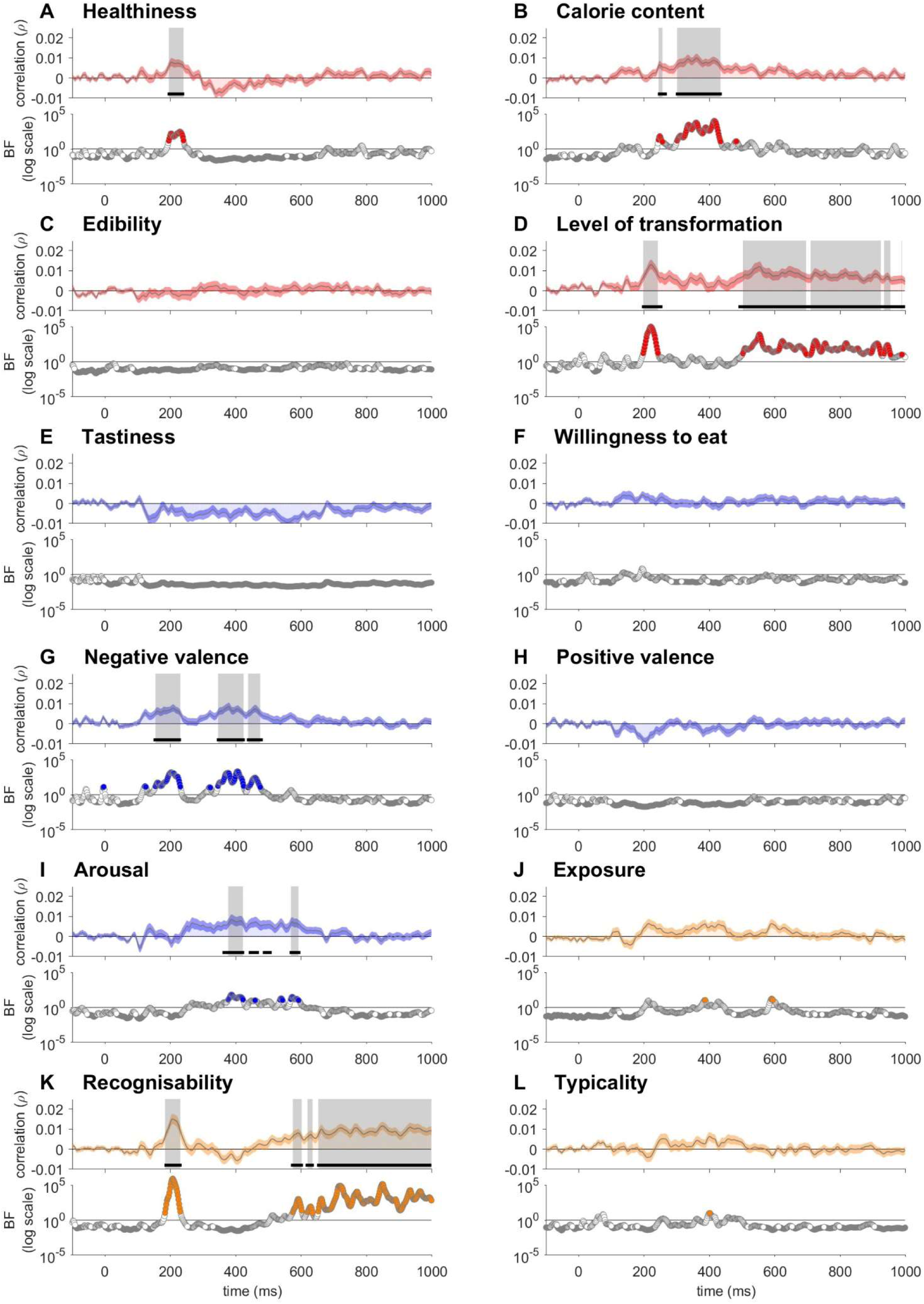
Correlations between the EEG and the food attributes while controlling for all other food attributes and visual features. Partial correlations between the food attribute RDMs and the neural RDMs, controlling for the other ll food attribute RDMs and the six visual feature RDMs. Nutritive properties (A-D) include healthiness, calorie content, edibility, and level of transformation. Hedonic properties (E-I) include tastiness, willingness to eat, negative valence, positive valence, and arousal. Familiarity attributes (J-L) include exposure, recognisability, and typicality. The shaded area around the plot lines shows the standard error of the mean. The horizontal black lines indicate time windows with statistically significant above-zero correlations after applying multiple comparisons corrections. The coloured circles indicate the time-points at which the BFs were greater than 10, while grey circles indicate the time-points at which the BFs were less than 1/10, and the white circles indicate the time-points at which the BFs were between 1/10 and 10. The grey shaded areas indicate the time windows at which the BFs were greater than 10 and there were statistically significant above-zero correlations after applying multiple comparisons corrections.

#### 3.3.2 Hedonic properties

Negative valence and arousal accounted for unique variance in the neural activity (Figure 6G, 6I). Negative valence was correlated from 154 ms, with an early peak at 209 ms, as well as subsequent windows of correlations from 34 ms. In contrast, arousal was correlated from 377 ms.

#### 3.3.3 Familiarity

Out of the familiarity attributes, only the recognisability model explained some unique variance in the EEG data (Figure 6K). There was an early peak at 207 ms and later, sustained windows of correlations starting from 574 ms and lasting until the end of the analysed time range.

## 4 Discussion

We comprehensively examined how patterns of EEG signals covary with food attribute judgements. Participants viewed food images while their EEG activity was recorded, and ratings on food attributes related to nutritive properties, hedonic properties, and familiarity were collected from a separate sample. Using RSA, we assessed the neural time-courses of each food attribute individually, while controlling for potential confounds associated with low- and mid-level visual features. We reveal that food attributes were represented in the neural activity rapidly and in parallel, in contrast to the idea that certain attributes are processed more quickly than others. Our findings suggest that decision-relevant features of foods may capture early attention and facilitate rapid appraisal processes. In light of the substantial covariation between the food attributes, we examined the broad underlying dimensions of foods using PCA and the unique contributions of individual attributes using partial correlation analyses. We found that the broad dimensions underlying individual attributes (how appetising and how processed the food is) were also represented in early neural signals. On the other hand, the neural activity also reflected information unique to a subset of individual attributes. These findings highlight that the structure and time-courses of food attribute representations can be mapped at multiple levels. Furthermore, we demonstrate the utility of examining the time-courses of neural representations in EEG activity by coupling a large EEG dataset and attribute ratings collected from a separate sample.

### 4.1 The neural time-courses of individual food attributes

We found that many food attributes shared similarities in their time-courses of correlation with the EEG signals. For many attributes, we observed an early peak in the correlation (∼200 ms from food image onset) as well as later, more sustained windows of encoding (from ∼400-650 ms). These findings are consistent with previous findings that some key food attributes are reflected in early (∼100-200 ms) and later, more sustained (∼300-700 ms) ERP components (Duraisingam et al., 2021; Harris et al., 2013; Meule et al., 2013; Rosenblatt et al., 2018; Schubert, Smith, et al., 2021; Toepel et al., 2009). Previous work has suggested that differences in ERP amplitudes evoked by foods may reflect differences in initial attention allocation (see review by Carbine et al., 2018). In line with this, early windows of encoding in neural activity (∼100-200 ms) may be reflective of early attentional processes evoked by foods that differ in decision-relevant features. On the other hand, activity related to stimulus value signals are thought to emerge later (∼400-500 ms from stimulus onset) in food decisions (Harris et al., 2013). We hypothesise that the later, sustained windows of encoding may reflect parallel appraisal of attributes before the integration of information towards a decision. While it has been well established that key food attributes (e.g., tastiness, healthiness, and calorie content) are reflected in the neural activity over these time windows when explicitly judging the foods on these attributes, we demonstrate that a wide array of attributes may be processed in parallel during food viewing. Taken together, salient features of foods may capture early attention, facilitating rapid valuation of the food on relevant attributes.

Researchers have speculated that activity related to decision-relevant food attributes occur in the ventromedial prefrontal cortex (vmPFC) in these temporally dissociable windows (Harris et al., 2011, 2013). The vmPFC is widely thought to be involved in the encoding of stimulus value across many different types of value-based decision-making (Hare et al., 2009; Kable & Glimcher, 2007; Rangel & Clithero, 2014). In dietary self-control literature, this later window (∼400-650 ms) is thought to be when stimulus value is modulated to exert self-control (e.g., increased weighting of healthiness; Harris et al., 2013). Our findings are in line with the idea that food attributes are represented and used by the brain for decision-making during these later time windows; however, our findings do not give us information about the specific brain regions involved in these processes.

Furthermore, we corroborate and extend on findings by Moerel et al. (2024) by applying a similar multivariate approach to map the neural time-courses of representations for a different, overlapping set of food attributes. Consistent with Moerel et al. (2024), we found that nutritive properties are represented in EEG activity earlier (∼100-350 ms) for level of transformation and calorie content and later (∼650 ms) for edibility. On the other hand, we found evidence for the representation of hedonic properties in the neural activity, while Moerel et al. (2024) did not find strong evidence for the representation of valence and arousal. We observed a later onset (∼600-650 ms) of the sustained window of encoding for hedonic properties, which may explain why Moerel et al. (2024) did not find strong evidence for the representation of these attributes at very rapid presentation rates (e.g., ∼100 ms per image). While numerous studies have shown that features of objects are represented in the EEG signals even at very rapid presentation rates (Contini et al., 2020; Grootswagers, Robinson, Shatek, et al., 2019; Marti & Dehaene, 2017; Mohsenzadeh et al., 2018), unlike early neural responses (<150 ms), later processing may be influenced by presentation rate (Grootswagers, Robinson, & Carlson, 2019). Representations at very rapid presentations rates (∼50 ms per image) were weaker and lasted for shorter durations compared to slower presentation rates (∼200 ms per image), potentially due to the increase in presentation duration leading to further consolidation of representations (Grootswagers, Robinson, & Carlson, 2019). Future studies may opt for an extended presentation duration (e.g., in the order of seconds) when investigating the appraisal of foods. Different stimulus sets used in the studies may be another contributing factor to this discrepancy. Selecting food stimuli to include edible but less appetising foods ensured that we were able to obtain sufficient variance in the hedonic properties. Achieving sufficient variance in the attributes of interest may be especially crucial for multivariate analyses that rely on fine-grained information to train a classifier or construct meaningful RDMs. Taken together, these findings shed light on the methodological differences (e.g., viewing duration, stimuli selection) that influence the mapping of the neural representations of different types of food attributes. Furthermore, we reveal the neural time-courses of a novel class of food attributes (familiarity), which appear to share similarities with the time-courses of hedonic properties. We add to a growing body of knowledge showing that key food features are rapidly represented in neural activity and highlight the importance of experimental design when assessing subtle patterns of neural activity.

Our findings do not support the idea that tastiness information is processed more quickly compared to healthiness information when viewing foods, revealing that the latter can in fact be represented in the EEG activity earlier than the former. This is consistent with some findings reported by Schubert, Rosenblatt, et al. (2021), where the decoding onset of healthiness ratings preceded the decoding onset of tastiness ratings when participants were not explicitly evaluating the foods on healthiness or tastiness. Categorising attributes in terms of their level of abstraction or short- or long-term relevance may pose a problem for food attributes, especially given the intercorrelated structure of these attributes. For example, calorie content can be conceptualised as an attribute with long-term relevance as it plays a role in nutrition, but also signals short-term tastiness due to its close relationship with the fat and sugar content of foods. For other attributes, such as typicality, recognisability, emotional arousal, or level of processing, their level of abstraction and relevance to short- or long-term goals is less clear and may be situational. In the context of a typical food cognition study that uses food images as stimuli rather than real foods, in which real food choice is not imminent, attributes may not differ so much in their short- or long-term relevance. When there is a real food choice at stake (e.g., when participants are aware that they will have to consume one of the foods that they choose in the experiment), we may observe that more immediately relevant attributes are indeed processed more quickly (e.g., Sullivan et al., 2015). Our results show that in typical lab settings, healthiness and other attributes that are more ambiguous in their level of abstraction can be processed rapidly. These findings lend support for the parallel encoding of multiple relevant attributes that inform a food choice.

### 4.2 Understanding food attribute covariation

We took a novel approach to address the issue of food attribute covariation, which historically has made it difficult to draw conclusions about the time-courses of correlations for individual food attributes. Previous studies have been limited in examining only one or a few food attributes at a time and proposing that any differences in neural activity reflected information about that food attribute without considering covarying attributes that may also contribute (Coricelli et al., 2019; Harris et al., 2013; Meule et al., 2013; Rosenblatt et al., 2018; Schacht et al., 2016; Schubert, Smith, et al., 2021; Stuldreher et al., 2023; Toepel et al., 2009). We collected ratings for a wide range of food attributes, which allowed us to explicitly investigate the structure of attribute covariation and conduct further analyses to examine the broader dimensions that may underlie individual attributes. In a complementary approach, we also assessed whether there were unique contributions from the individual attributes not shared with the other attributes.

We identified that the 12 food attribute ratings were clustered into a few groups that were strongly positively correlated (Figure 2B). The largest cluster included familiarity attributes (exposure, recognisability, and typicality) and many of the hedonic properties (tastiness, willingness to eat, negative and positive valence). The second largest cluster included many nutritive properties (healthiness, calorie content, and level of transformation). Arousal and edibility were each found to be relatively distinct from the other attributes. Consistent with this, the time-courses of correlations with the EEG signals were similar within the clusters (Figure 5), indicating that the covariation among food attributes were reflected in the neural signals over time. This suggests that shared underlying dimensions may be driving the covariation between the food attributes, and these dimensions may be encoded in the neural activity.

Using PCA, we demonstrated that two broad underlying dimensions can capture most of the variance in multiple correlated attributes. We further showed that these dimensions display two distinct time-courses of representations. The first dimension, reflective of how appetising the food is, appears to be processed in two stages, with an early brief window (∼200 ms) followed by a later, more sustained window (∼600 ms and onwards). This is consistent with findings by Carbine et al. (2018) that reported that foods, in comparison to neutral cues or less palatable foods, elicit greater ERP amplitudes that reflect early attentional allocation (e.g., increased P2). The early window in our results may reflect early attentional allocation toward foods that are more appetising, while the later window may correspond to higher-level appraisal processes. In contrast, the second dimension, reflective of how processed the food is, appears to be represented in a sustained manner from ∼150 ms onwards. We found distinct time-courses of representations for the two dimensions, most notably in the onset time for the sustained window of encoding. One potential explanation for this difference may be that appraising foods on how appetising it is involves cognitive processes such as memory retrieval of past eating experiences, interoception (e.g., evaluation of one’s current hunger level), or mental simulation, while appraising foods on how processed it is may not involve such processes to the same degree.

We demonstrate a novel way to get closer to the dimensions most salient for the brain, using principal components rather than individual attributes. The brain may map representations of foods on dimensions that correlate with, but are not exactly, the attributes we choose for our studies. On the other hand, we also show that half of the food attributes (healthiness, calorie content, level of transformation, negative valence, arousal, and recognisability) explained some amount of unique variance in the neural data not shared with any other attribute. In particular, level of transformation and recognisability explained unique variance in later, more sustained time windows. Our approach highlights the multilevel structure of food representations, revealing that how appetising and how processed the food is may broadly capture many decision-relevant attributes and are represented by the brain, while at the same time, the neural signals also contain information uniquely relevant to certain individual attributes.

Future work examining the neural correlates of food attribute processing should carefully consider possible multilevel, correlated structures of food attribute representations. For example, attribute ratings may be used to derive a food stimulus set with minimal correlations across targeted sets of attributes, or decorrelated stimulus sets may be curated for each individual participant. Alternatively, food attribute ratings could be collected first before selecting a subset of food stimuli that are not strongly correlated on relevant attributes to investigate the neural correlates of specific food attributes. Our approach using PCA may be particularly useful for identifying and mapping patterns of shared variance across food attributes.

### 4.3 A novel approach combining EEG recordings with online food attribute ratings

We demonstrate a novel approach for examining food representations in the human brain. We showed that attribute ratings collected from a separate sample can be used in conjunction with EEG data to examine the neural time-courses of representations. Future studies may consider using available datasets containing EEG data recorded during food viewing to examine representations of foods. For example, future work can involve collecting ratings on novel food attributes from a separate sample and applying RSA to examine whether these novel attributes are represented in the EEG signals in an available dataset. As EEG data acquisition can be time-consuming and expensive, this approach provides a potentially more efficient way to study the neural processing of foods.

### 4.4 Conclusion

In this study, we coupled food attribute rating data with a large EEG dataset. We delineate the neural time-courses of food attribute representations at different levels, from individual attributes to broad underlying dimensions. Our findings suggest that salient food features may capture early attention, facilitating rapid and parallel appraisal of foods on multiple relevant attributes over similar time-courses. Beyond the neural time-courses of individual attributes, we reveal that broad underlying dimensions of how appetising and how processed the food is are also represented in the neural activity with distinct time-courses. Future studies may leverage available EEG datasets to explore the neural correlates of novel food attribute processing or consider alternative methods for parsing out representations of highly covarying attributes.

## 5 CRediT authorship contribution statement

Violet J. Chae: Conceptualisation, Data curation, Formal analysis, Writing – original draft, Writing – review and editing.

Tijl Grootswagers: Methodology, Supervision, Conceptualisation, Writing – review and editing.

Stefan Bode: Conceptualisation, Supervision, Funding acquisition, Methodology, Writing – review and editing.

Daniel Feuerriegel: Project administration, Supervision, Conceptualisation, Methodology, Writing – review and editing.

## 6 Data and code availability

The data and analysis code can be found via the Open Science Framework (https://osf.io/w8c9d/) at the time of publication.

## 8 Acknowledgement of funding

This project was supported by an Australian Research Council Discovery Early Career Researcher Award to D.F. (ARC DE220101508), an Australian Research Council Discovery Early Career Researcher Award to T.G. (ARC DE230100380), and a Research Training Program scholarship awarded to V.J.C. from the Australian Government. Funding sources had no role in study design, data collection, analysis or interpretation of results.

## 9 Ethical statement

The experiment was approved by the Human Research Ethics Committee (ID 24850) of the University of Melbourne.

