## Supplementary Material for "Characterising the neural time-courses of food attribute representations"

Redmond Barry Building,

University of Melbourne, VIC 3010, Australia

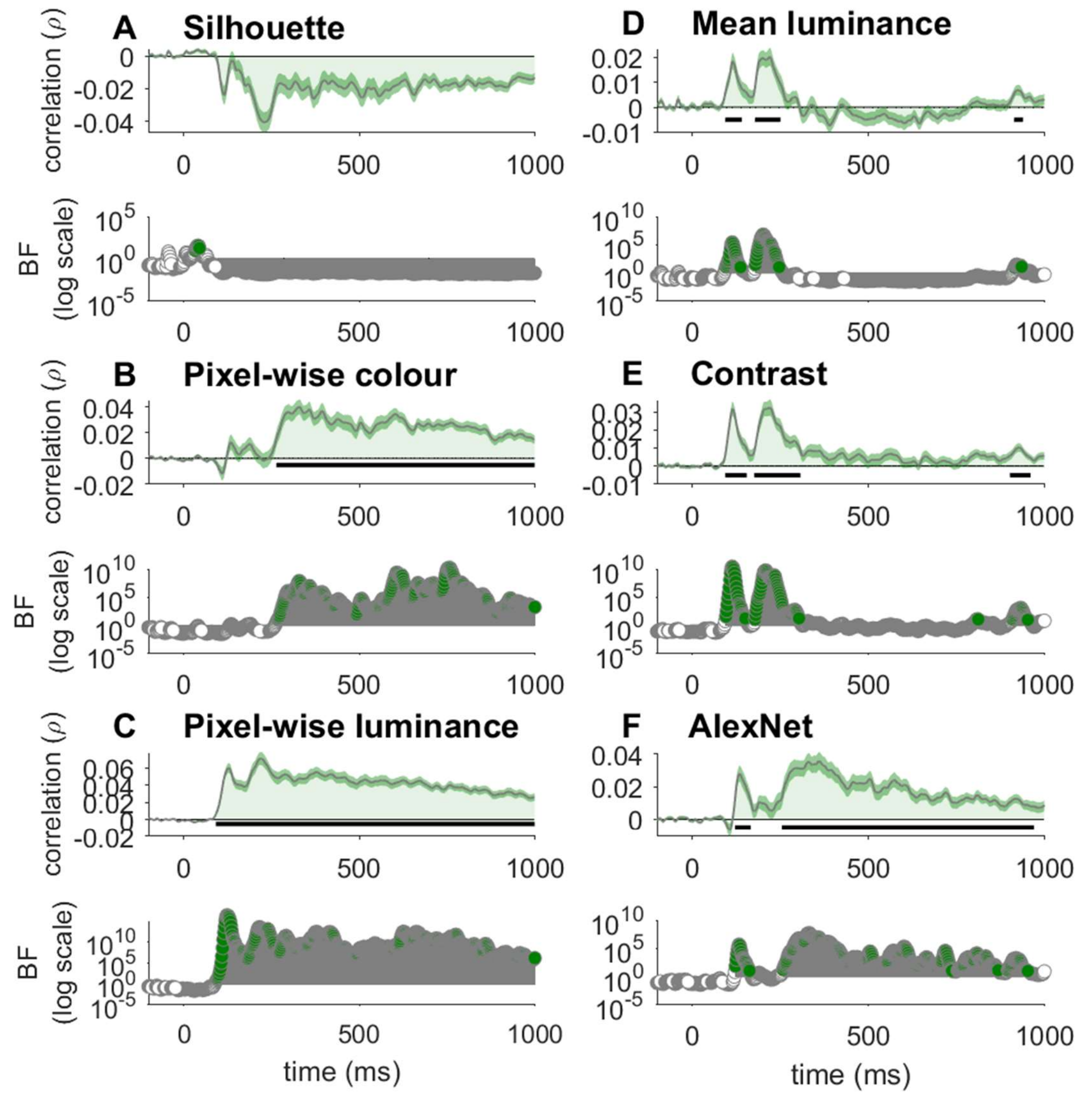


**Supplementary Figure 1. Correlations between EEG signals and low- and mid-level visual features.** Correlations between the silhouette (A), pixel-wise colour (B), pixel-wise luminance (C), mean luminance (D), contrast (E), and AlexNet (F) models and the neural RDMs. The shaded area around the plot lines shows the standard error of the mean. The coloured circles indicate the time-points at which the BFs were greater than 10. The horizontal black lines indicate time windows with statistically significant above-zero correlations after applying multiple comparisons corrections.
